# Circadian Gene Expression in Mouse Renal Proximal Tubule

**DOI:** 10.1101/2022.08.26.505418

**Authors:** Molly A. Bingham, Kim Neijman, Hiroaki Kikuchi, Hyun Jun Jung, Brian G. Poll, Viswanathan Raghuram, Euijung Park, Chin-Rang Yang, Chung-Lin Chou, Lihe Chen, Jens Leipziger, Mark A. Knepper, Margo Dona

**Affiliations:** Epithelial Systems Biology Laboratory, Systems Biology Center, National Heart, Lung, and Blood Institute, National Institutes of Health, Bethesda, Maryland; Department of Internal Medicine, Radboud University Medical Center, Nijmegen, the Netherlands; Division of Nephrology, Department of Medicine, Johns Hopkins University School of Medicine, Baltimore, Maryland; Department of Biomedicine, Physiology, Aarhus University, 8000 Aarhus, Denmark

**Keywords:** Circadian variation, proximal tubule, transcriptomics

## Abstract

Circadian variability in kidney function has long been recognized but is often ignored as a potential confounding variable in *in vivo* physiological experiments. To provide a guide for physiological studies on the kidney proximal tubule, we have now created a data resource consisting of expression levels for all measurable mRNA transcripts in microdissected proximal tubule segments from mice as a function of the time of day. This approach employs small-sample RNA-sequencing (RNA-seq) applied to microdissected renal proximal tubules including both S1 proximal convoluted tubules (PCTs) and S2 proximal straight tubules (PSTs). The data were analyzed using *JTK-Cycle* to detect periodicity. The data are provided as a user-friendly web page at https://esbl.nhlbi.nih.gov/Databases/Circadian-Prox/. In PCTs, 234 transcripts were found to vary in a circadian manner (3.7 % of total quantified). In PSTs, 334 transcripts were found to vary in a circadian manner (5.3 % of total quantified). Transcripts previously known to be associated with corticosteroid action and transcripts associated with increased flow were found to be overrepresented among circadian transcripts peaking during the “dark” portion of the day (Zeitgeber 14-22), corresponding to the peak levels of corticosterone and glomerular filtration rate in mice.

**Blurb:** Circadian variation in gene expression can be an important determinant in the regulation of kidney function. The authors used RNA-seq in microdissected proximal S1 and S2 segments to identify transcripts that vary in a circadian manner. The data were used to construct a user-friendly web resource.

## INTRODUCTION

The design of any *in vivo* physiological experiment addressing kidney function is challenging. Aside from the experimental variable of interest, there are many potential confounding variables that could influence the interpretation of results. For example, the sex of the animal can have an important influence on certain physiological functions and gene expression (1-11). Other important variables include the age of the animals under study, diet, and the time of day when the observations are made. Circadian variability in kidney function has long been recognized (12-16), but is often ignored in modern physiological experiments. However, recent studies have reported significant circadian variability in gene expression in the kidney (17-26). Circadian variation in renal gene expression can occur either through cell-autonomous behavior involving the canonical circadian signaling network resident in kidney epithelial cells or through responses to circadian variation of external signals such as hormone levels, renal nerve activity or hemodynamics (27-31). Two external signals of particular interest are circadian changes in glomerular filtration rate (GFR) (32, 33) that produce changes in flow rate in the proximal tubule, and circadian changes in adrenal corticosteroids in blood plasma (34, 35) that can result in changes in gene expression through the ligand-activated transcription factors Nr3c1 and Nr3c2. In rodents, GFR and corticosteroid levels are highest during the active, lights-out period (ZT>12) (32-35).

To provide a guide for physiological studies on the kidney proximal tubule, we have now created a data resource consisting of expression levels for all measurable mRNA transcripts in microdissected proximal tubule segments from mice as a function of the time of day. This approach employs small-sample RNA-sequencing (RNA-seq) applied to microdissected renal proximal tubules (36), including both S1 proximal convoluted tubules and S2 proximal straight tubules. The data are provided as a user-friendly web page at https://esbl.nhlbi.nih.gov/Databases/Circadian-Prox/.

Although the main goal of this study is to provide a reference data set for use in experimental design, it should be possible to answer some simple questions with the new data. First, what fraction of transcripts that are expressed in the proximal tubule of the mouse kidney undergo circadian variation? Second, is the variation seen mainly due to cell autonomous behavior via recognized circadian signaling pathways intrinsic to proximal tubule cells, or is the variation due to circadian variations in external stimuli? Regarding the latter question, we analyze the genes that show circadian variability to determine whether known flow-dependent or corticosteroid-dependent transcripts vary in a manner related to circadian variability in GFR and plasma corticosteroid levels.

## METHODS

### Animals

The study was done in mice (male C57BL/6, 4-6 weeks) maintained in an environmentally controlled animal room adjusted to a 12hr:12hr light:dark cycle and were given ab libitum access to water and food in the animal facility of the Department of Biomedicine, Aarhus University (Removal of mouse organ tissue after sacrificing the animal complies with the Danish animal welfare regulations and the general license for animal caretaking at *Aarhus Biomedicine* is granted by the Animals Experiments Inspectorate (2021-15-0206-00018/BES). Three animals were selected at each of 6 time points starting at 8 a.m. (lights on, Zeitgeber 0) and then at every-four-hour junctures. Each of the three animals at each time point were studied on separate days. The mice were rapidly euthanized by cervical dislocation.

### Microdissection of S1 and S2 proximal tubules

Transverse sections of the central part of the kidney containing the entire corticomedullary axis were made as described by Wright and Knepper (37). The sections were put in dissection buffer (NaCl [120 mM], Na_2_HPO_4_ [2.5 mM], sodium acetate, [5 mM], MgSO_4_ [1.2 mM], KCI [5 mM], CaCl_2_ [2 mM], glucose [5.5 mM], HEPES [5 mM], pH 7.4) freshly supplemented with alanine [6 mM], trisodium citrate [1 mM], heptanoate [1 mM] and glycine [4 mM] in DNase I and RNase free water). Using a Wild M8 dissection stereomicroscope (Wild Heerbrugg, Heerbrugg, Switzerland), S1 proximal convoluted tubules were dissected from the cortical labyrinth and S2 proximal straight tubules were dissected from cortical medullary rays. The S1 segments were distinguished by their attachments to glomeruli. The S2 segments originated from the outer cortex. Each time point was represented by 3 samples for both S1 and S2 segments, each from a different mouse. The single tubules were then washed twice in PBS before being transferred to Trizol. The entire procedure was carried out on ice or on cooled surfaces.

### RNA Extraction and RNA-seq

For the isolation of RNA from the microdissected proximal tubules, 0.2 ml chloroform was added per ml of Trizol. After vigorously shaking for 20 seconds, the mixture was incubated at room temperature for 2-3 minutes. After centrifugation (10,000x g, 18 minutes, 4°C), the aqueous phase was removed and an equal volume of 100% EtOH was added. The mixture was then loaded on an RNeasy column (Qiagen RNeasy micro kit) and the manufacturers’ protocol, including the DNase I treatment, was followed. RNA-seq was performed at GenomeScan B.V., Leiden, The Netherlands. The NEBNext Single Cell/Low Input RNA library Prep kit for Illumina was used to process the samples. The sample preparation was performed according to the protocol “NEBNext Single Cell/Low Input RNA Library Prep kit for Illumina”(NEB #E6420S/L). Briefly, cDNA template is generated from the mRNA. The template is fragmented and sequencing adapters with unique indexes are ligated, the ligated product is amplified with a PCR. The quality and yield after sample preparation was measured with the Fragment Analyzer. The size of the resulting products was consistent with the expected size distribution (peak fragment size between 200-500 bp). Clustering and DNA sequencing using the NovaSeq6000 was performed according to manufacturer’s protocols. 1.1 nM of cDNA was used. NovaSeq control software NCS v1.6 was used in demultiplexing and generation of fastq files.

### Identification of circadian transcripts in proximal tubules

Transcripts per million (TPM) values were calculated for each sample. Full RNA-seq data is archived at (https://doi.org/10.25444/nhlbi.20533536). Thedata were filtered to include only transcripts with a maximum TPM>7 to remove noisy, low abundance mRNAs. The data were analyzed using *JTK-Cycle* (38) to detect periodicity (see https://doi.org/10.25444/nhlbi.20533536 for calculations). For this analysis, circadian genes were those with a period 20 or 24 hours, and a curve fit significance level of P<0.05 by nonparametric test described by Hughes et al (38). Analyzed data were shared via user-friendly web pages accessible at https://esbl.nhlbi.nih.gov/Databases/Circadian-Prox/.

### Identification of flow-dependent transcripts

To create a gene set for transcripts up- or down-regulated by changes in flow in renal epithelia, we used data downloaded from a prior study (39) in which the investigators grew mouse IMCD cells in a flow chamber and varied flow rate. TPM values were compared between cells in a static environment and cells subjected to flow. For the analysis, “Flow TPM” was divided by “Static TPM”. Responses to flow were not reported directly in the original paper and we now report these data to readers as an additional resource at https://esbl.nhlbi.nih.gov/Databases/IMCD3_flow_effect/RD.htm. 208 transcripts were identified as increased (log_2_[Flow/Static] > 1; P < 0.05; “Flow UP” gene list). Similarly, 140 transcripts were found to be decreased (log_2_[Flow/Static] < -1; P < 0.05; “Flow DOWN” gene list). The standard deviation (SD) for log_2_ (static/static) averages 0.34, indicating that 99 percent of the intrinsic variability of the method would be [-0.88, +0.88], calculated as ±2.576*SD. Thus, the criterion used (|log_2_[Flow/Static]|>1) approximates the 99 percentile criterion and can be considered very stringent. The joint probability of a false positive (calculated from P and the 99% confidence interval) would be 0.05 X (1-0.99) = 0.0005.

### Identification of corticosteroid-dependent transcripts

We have used gene lists from several sources to create a “corticosteroid-induced” gene list (40-47). The list is the union of the lists curated at https://esbl.nhlbi.nih.gov/Signaling-Pathways/Glucocorticoid-Targets/ and at https://esbl.nhlbi.nih.gov/Signaling-Pathways/Aldo-Targets/.

### Bioinformatics

Gene Ontology (GO) term enrichment analysis of circadian transcript lists was performed using the *Database for Annotation, Visualization and Integrated Discovery* (DAVID; https://david.ncifcrf.gov/) with the full list of transcripts quantified serving as a reference background (48). Chi-square analysis was carried out using an online calculator (https://www.socscistatistics.com/tests/chisquare2/default2.aspx).

## RESULTS AND DISCUSSION

We carried out RNA-Seq transcriptomic analysis of microdissected proximal convoluted tubules (PCT, S1 segment) and proximal straight tubules (PST, S2 segment) from mice at 4-hour intervals starting at Zeitgeber 0 (lights on 8 a.m.). The data were analyzed using *JTK-Cycle* (38). The data were used to create an online resource, which allows searching, browsing or download of data (https://esbl.nhlbi.nih.gov/Databases/Circadian-Prox/). In PCTs, 6296 transcripts were quantified and 234 of these were found to vary in a circadian manner (3.7 %) (period 20 or 24 hours, P<0.05 by nonparametric test described by Hughes et al (38)). In PSTs, 6304 transcripts were quantified and 334 of these were found to vary in a circadian manner (5.3 %) (period 20 or 24 hours, P<0.05). 35 transcripts showed significant circadian variation in both PCT and PST including *Heat Shock Protein 90 Alpha Family Class B Member 1* (Hsp90ab1), *N-Myc Downstream Regulated 1* (Ndrg1), *Oxidative Stress Induced Growth Inhibitor 1* (Osgin1), *Phosphoenolpyruvate Carboxykinase 1* (Pck1), *RAR Related Orphan Receptor C* (Rorc), and *Serum/Glucocorticoid Regulated Kinase 1* (Sgk1). Web users can visualize the graphed time-course data by hovering over each official gene symbol at https://esbl.nhlbi.nih.gov/Databases/Circadian-Prox/Circadian_Amplitude_PCT.html for PCT circadian transcripts and at https://esbl.nhlbi.nih.gov/Databases/Circadian-Prox/Circadian_Amplitude_PST.html for PST circadian transcripts. **Figures 1 and 2** show examples of circadian transcripts that peak during the lights-on period ZT0-ZT12 (A) and the “lights-off” period ZT12-ZT24 (B) for PCT and PST, respectively.

**Figure 1:**
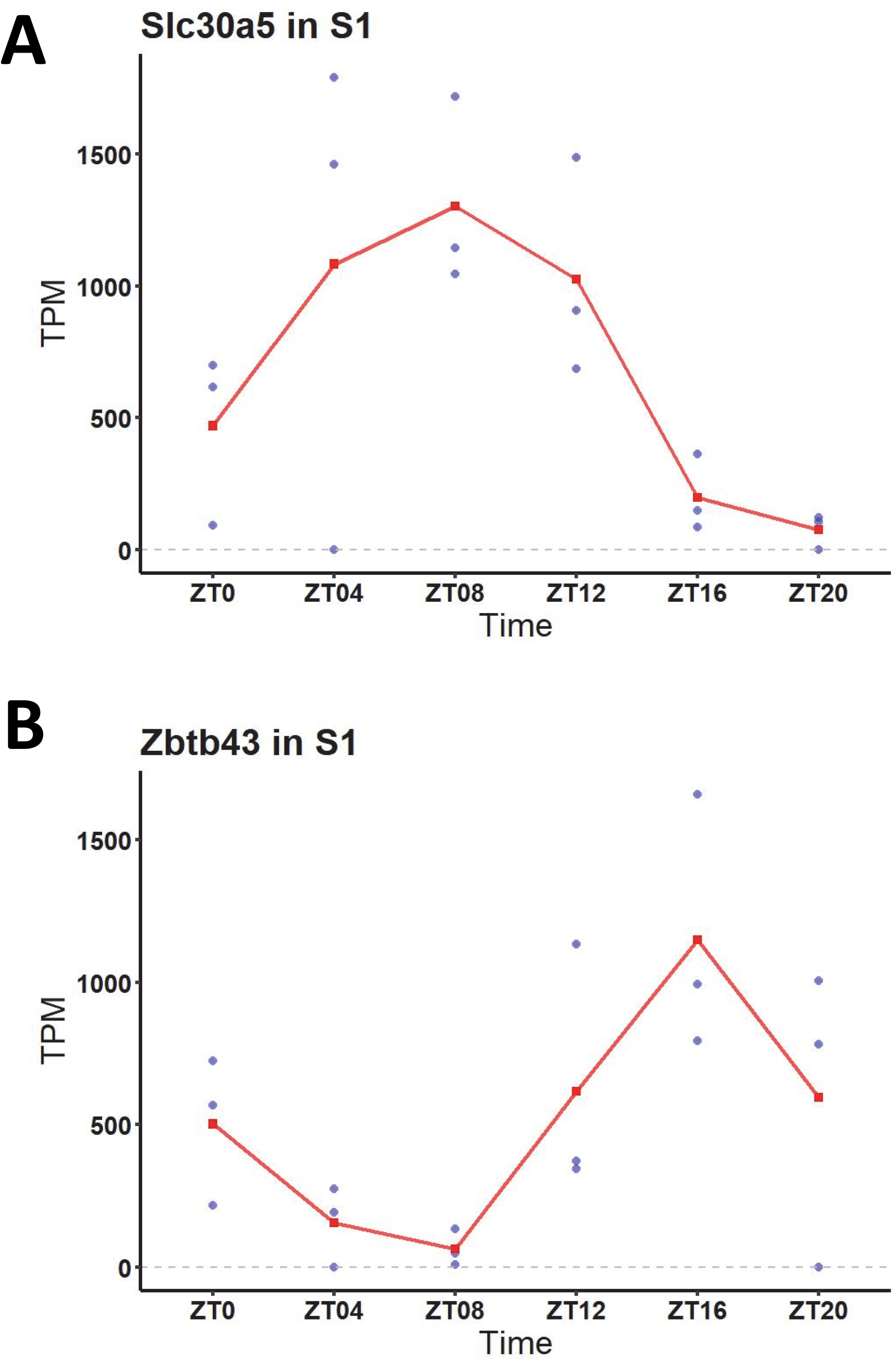
Two examples of time course data for transcripts using images downloaded from the data-sharing web page at https://esbl.nhlbi.nih.gov/Databases/Circadian-Prox/Circadian_Amplitude_PCT.html. Transcripts are indicated by official gene symbols. “S1” refers to the PCT. “ZT” is short for Zeitgeber, representing the time point in hours since lights-on time. **A**. Transcript coding for *solute carrier family 30* (zinc transporter) *member 5* (Slc30a5) in the PCT has a phase of 8 hours, corresponding to a peak amplitude 8 hours after lights on. **B**. The transcript coding for *zinc finger and BTB domain containing 43* (Zbtb43) in the PCT has a phase of 18 hours (peak amplitude approximately 6 hours after lights off).

**Figure 2:**
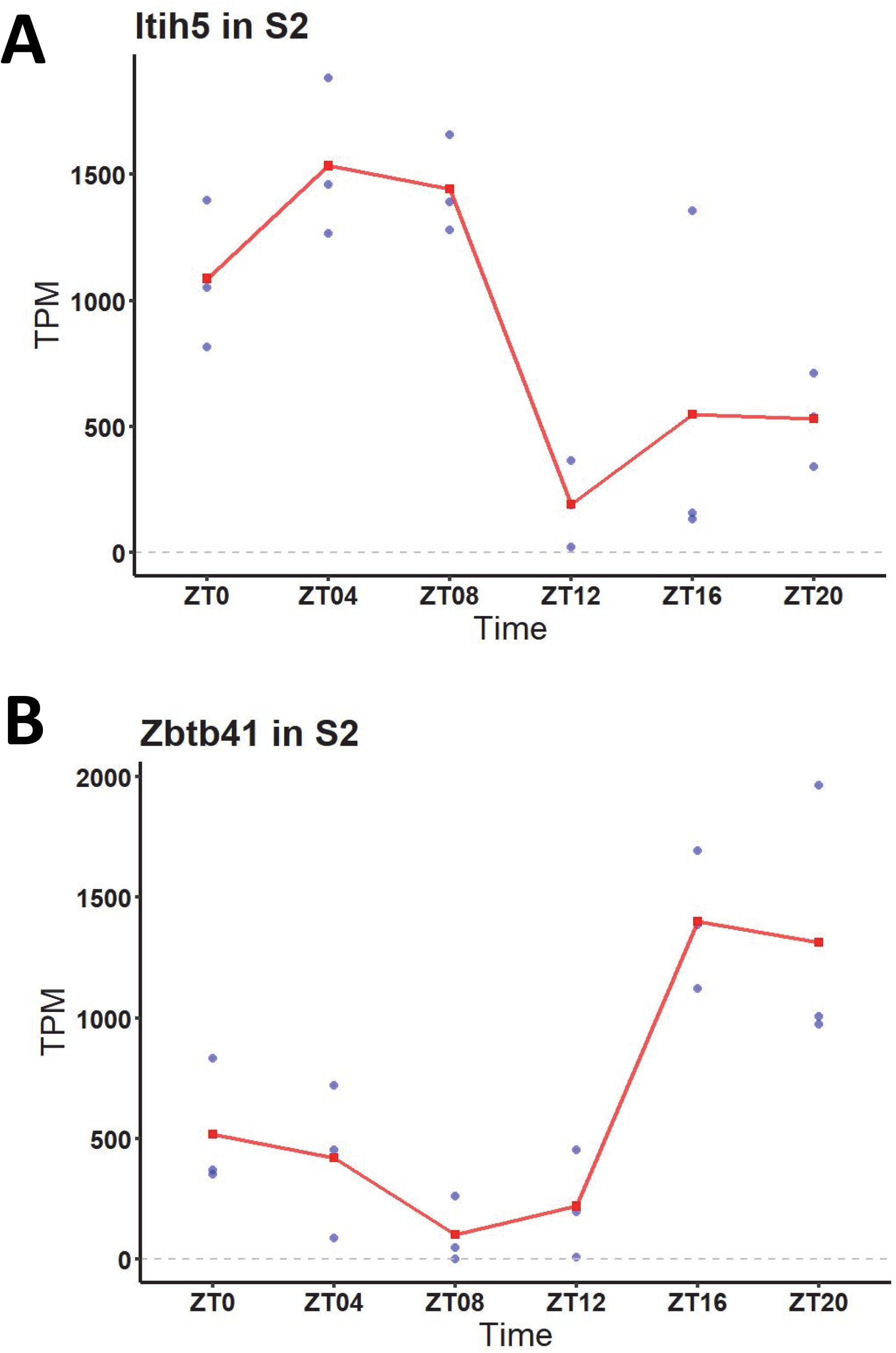
Two examples of time course data for transcripts using images downloaded from the data-sharing web page at https://esbl.nhlbi.nih.gov/Databases/Circadian-Prox/Circadian_Amplitude_PST.html. Transcripts are indicated by official gene symbols. “S2” refers to the PST. “ZT”, Zeitgeber, representing the time point in hours since lights-on moment. **A**. Transcript coding for *inter-alpha-trypsin inhibitor heavy chain H5* (Itih5) in the PST has a phase of 4 hours, with a peak amplitude about 4 hours after lights on. **B**. The transcript coding for *zinc finger and BTB domain containing 41* (Zbtb41) in the PCT has a phase of 20 hours, peaking approximately 8 hours after lights off.

As summarized in the *Introduction*, circadian behavior in a particular cell type can be intrinsic (cell autonomous) or imposed by outside signaling factors (29). The canonical intrinsic circadian transcripts are *Aryl Hydrocarbon Receptor Nuclear Translocator Like* (Arntl or Bmal1), *Clock Circadian Receptor* (Clock), *Periodic Circadian Regulator 1, 2 and 3* (Per1, Per2, Per3), *Timeless Circadian Regulator* (Timeless), *Cryptochrome Circadian Regulator 1 and* 2 (Cry1, Cry2), *RAR Related Orphan Receptor A, B and C* (Rora, Rorb, Rorc), and *Nuclear Receptor Subfamily 1 Group D Member 1 and 2* (Nr1d1, Nr1d2) (29). Among these, only Rorc showed circadian variation of transcript abundance with peak expression at Zeitgeber 20, https://esbl.nhlbi.nih.gov/Databases/Circadian-Prox/). Note, however, prior studies have provided strong evidence for important roles for components of the classical circadian signaling pathway in regulation of various aspects of kidney function (17-26, 49).

To address what extrinsic signals may have been responsible for the circadian behavior seen in this study, we carried out pathway analysis to identify possible upstream signals that may have contributed to circadian gene expression in PCT (**Table 1** (50-52)) and PST (**Table 2** (50-52)). To do this, we identified circadian transcripts that peaked either during the lights-on period (Zeitgeber 2-10) or the lights-off period (Zeitgeber 14-22), combining the lists for PCT and PST. We asked whether we could identify pathways in which gene sets are over-represented (“enriched”) among lights-on or lights-off circadian transcripts relative to the entire set of expressed transcripts. To do this, we employed existing curated datasets from standard sources (GSEA Hallmark, KEGG). However, no appropriate gene sets were identifiable to cover the two hypotheses made in the *Introduction* for the possible origin of extrinsic circadian behavior: 1) that gene expression changes over the course of the day are due to variations in flow resulting from circadian variation in glomerular filtration rate (GFR); or 2) that gene expression changes over the course of the day are due to variations in corticosteroid levels, with signaling through the glucocorticoid receptor (Nr3c1) or the mineralocorticoid receptor (Nr3c2). Accordingly, we curated two data sets to cover these hypotheses (**Methods**). The pathways in which gene sets are enriched among circadian transcripts that peak during the lights-on period relative to the entire set of expressed transcripts are shown in **Table 1**. The only pathway that was significantly enriched was “p53 signaling”. The pathways in which gene sets are enriched among circadian transcripts that peak during the lights-off period relative to the entire set of expressed transcripts are shown in **Table 2**. The two pathways that were significantly enriched are “Corticosteroid-induced” and “Flow UP”. These two pathways peak at a time similar to or slightly delayed from the peak in GFR and proximal flow (32, 33) and corticosterone levels (34, 35).

**Table 1.**
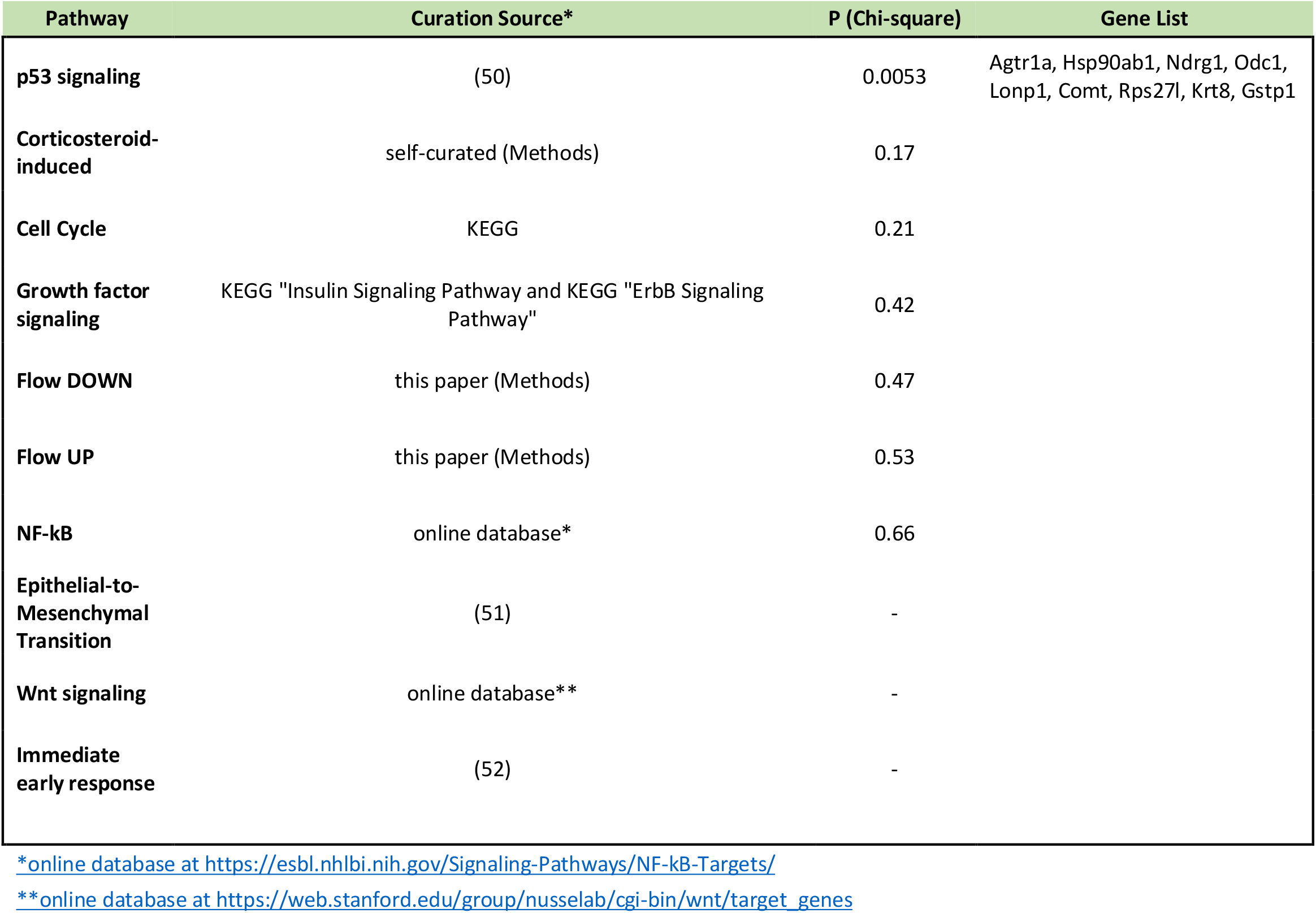
Pathways enriched or non-enriched in circadian genes peaking during ‘lights on’ (ZT 2-10)

**Table 2.** Pathways enriched or non-enriched in circadian genes peaking during ‘lights off’ (ZT 14-22)

| Pathway | Curation Source* | P (Chi-square) | Gene List |
| --- | --- | --- | --- |
| **Corticosteroid-induced** | self-curated (Methods) | <0.00001 | Tsc22d3, Sgk1, Usp2, Angptl4, Ddx5, Hsp90ab1, Mknk2, Pck1 |
| **Flow UP** | this paper (Methods) | 0.0013 | Ubc, Osgin1, Sgk1, Slc25a25, Angptl4 |
| **Flow DOWN** | this paper (Methods) | 0.063 |  |
| **Immediate early response** | (52) | 0.16 |  |
| **Growth factor signaling** | KEGG "Insulin Signaling Pathway and KEGG "ErbB Signaling Pathway" | 0.21 |  |
| **Wnt signaling** | online database** | 0.46 |  |
| **p53 signaling** | (50) | 0.83 |  |
| **Cell Cycle** | KEGG | 0.96 |  |
| **Epithelial-to-Mesenchymal Transition** | (51) | - |  |
| **NF-kB** | online database* | - |  |

\*online database at https://esbl.nhlbi.nih.gov/Signaling-Pathways/NF-kB-Targets/
\*\*online database (Wnt signaling) from: https://web.stanford.edu/group/nusselab/cgi-bin/wnt/target_genes

### Gene Ontology Term Analysis

To identify biological processes associated with the circadian genes, we used DAVID (Database for Annotation, Visualization and Integrated Discovery, NIAID) to identify *Gene Ontology Biological Process* terms enriched in the lights-on and lights-off period transcripts (**Table 3 and 4**, respectively). Common themes among biological processes enriched among transcripts peaking in the lights-on period were “ATP synthesis”, “Acute inflammatory response”, “Water homeostasis”, “DNA integrity” and “Translation”. Common themes among biological processes enriched among lights-off genes were “Glutathione metabolism”, “Apoptosis”, and “Regulation of transcription”. Given the large number of transcripts in the “Regulation of transcription” category, we asked what transcription factor transcripts vary in a circadian manner.

**Table 3.** *Gene Ontology Biological Process* terms significantly enriched in circadian genes that peak during ‘lights on’ period

| Gene Ontology Biological Process Term | Fold Enrichment | P (Fisher Exact) | Genes |
| --- | --- | --- | --- |
| ATP synthesis coupled electron transport | 3.1 | 0.0021 | mt-Nd6, Chchd2, mt-Nd5, mt-Co1, Ndufs2, Coq9, mt-Nd2, Cox5b, Cox6a1 |
| Acute inflammatory response | 4.6 | 0.0038 | Plscr1, Gstp1, Ogg1, Serpinf2, Fcgr2b |
| Water homeostasis | 4.2 | 0.013 | Elovl1, Stard7, Trpv4, Stmn1 |
| DNA integrity checkpoint | 2.9 | 0.027 | Syf2, Rps27l, Cdc5l, Rpl26, Nae1 |
| Translation | 1.5 | 0.031 | Mrps16, Mrps31, Rps27l, Mrps30, Rpl8, Rpl9, Mrpl10, Ptms, Tmed2, Rpl13, Rwdd1, Cnot6l, Mrpl49, Cirbp, Tma7, Mrpl28, Mrps21, Rpsa, Cnot10, Rps25, Eif5, Kars, Cdk4, Rpl26, Eif3d, Eif3b |

**Table 4.** Gene Ontology Biological Process terms significantly enriched in circadian genes that peak during ‘lights off’ period

| Gene Ontology Biological Process Term | Fold Enrichment | P (Fisher Exact) | Genes |
| --- | --- | --- | --- |
| Glutathione metabolic process | 5.1 | 0.0027 | Nat8f6, Nat8f5, Nat8, Oplah, Nat8f7 |
| Apoptotic process | 1.7 | 0.0098 | Prnp, Ddx5, Hsp90ab1, Tsc22d3, Rorc, Osgin1, Bfar, Mff, Ccar1, Herpud1, Dnaja1, Nat8f6, Nat8, Nuak2, Nat8f5, Mknk2, Nbn, Sgk1, Nat8f7, Tnfrsf21, Dedd, Nck1 |
| Regulation of transcription, DNA-templated | 1.4 | 0.039 | Prnp, Ddx5, Cbx3, Tsc22d3, Usp2, Rorc, Zbtb43, Tcf20, Bfar, Gtf2h3, Baz1b, Klf15, Chd1, Phc3, Zbtb41, Trak2, Ccar1, Pcgf1, Ccnl2, Pck1, Dedd, Gm45871, Cggbp1, Nck1 |
| Regulation of gene expression | 1.3 | 0.048 | Ddx5, Pcx, Zbtb43, Rorc, Tcf20, Chd1, Zbtb41, Trak2, Ccar1, Nat8f6, Nat8f5, Mknk2, Pcgf1, Ccnl2, Pck1, Nat8f7, Dedd, Nck1, Prnp, Cbx3, Tsc22d3, Usp2, Bfar, Gtf2h3, Baz1b, Klf15, Phc3, Clk1, Nat8, Tardbp, Cggbp1, Gm45871 |

### Transcription Factors

Gene regulatory networks typically involve multiple transcription factors. **Table 5** (53-63) shows the 13 transcription factors that vary significantly in a circadian manner. Many of these either have already been demonstrated to vary in a circadian manner in other cell types or have been associated with circadian behavior of other gene (53-63). One example is the transcription factor *MAF BZIP Transcription Factor* (Maf), which peaks in daytime and, according to GSEA Hallmark Signature data set MAF_Q6, maps to several genes seen to vary in a circadian manner that peak in the daytime, viz. *8-Oxoguanine DNA Glycosylase* (Ogg1), *HNF1 Homeobox B* (Hnf1b), *Cell Division and Cell Cycle Like* (Cdc5l), *Mitochondrial Ribosomal Protein L49* (Mrpl49), and *Adipogenesis Associated Mth938 Domain Containing* (Aamdc). Further study of these 13 transcription factors in the proximal tubule seems warranted.

**Table 5.** Transcription factors among proximal tubule circadian genes

| Gene | Peak Time | Period (hours) | Phase (hours) | Prior Evidence |
| --- | --- | --- | --- | --- |
| **Glis3** | Lights on | 20 | 4 | (53) |
| **Cdc5l** | Lights on | 24 | 4 | - |
| **Maf** | Lights on | 24 | 4 | - |
| **Mlx** | Lights on | 24 | 6 | (54-56) |
| **Hnf1b** | Lights on | 20 | 6 | (57) |
| **Nr1h4** | Lights on | 20 | 8 | (58) |
| **Klf9** | Lights on | 24 | 10 | (59) |
| **Klf15** | Dark | 20 | 14 | (60) |
| **Cers2** | Dark | 24 | 16 | (61) |
| **Zbtb43** | Dark | 24 | 18 | - |
| **Tsc22d3** | Dark | 24 | 18 | (62) |
| **Rorc** | Dark | 24 | 20 | (63) |
| **Zbtb41** | Dark | 24 | 20 | - |

Since the writing of this manuscript, a preprint publication has appeared reporting RNA-seq in whole kidney samples from mice sampled at 4-hour intervals throughout the day [Wolff et al. doi: https://doi.org/10.1101/2022.04.27.489594]. Although that study was not specifically focused on proximal tubule cells, we note that about 65% of the total protein content of the epithelial cells of the kidney are contributed by proximal tubule cells (64). Thus, the whole kidney transcriptomic data of Wolff et al could be expected to be chiefly a reflection of gene expression in proximal tubule, justifying a comparison between the two studies. Of the 25 protein-coding transcripts that show strong circadian behavior in both S1 and S2 proximal tubules in our study, 20 showed significant circadian behavior in the whole kidney study of Wolff et al. based on their use of the *JTK-cycle* program (criterion: P<0.05 for sine wave regression, see **Supplemental Figure 1** at https://doi.org/10.25444/nhlbi.20533536.). In contrast, in a randomly selected set of 25 protein-coding transcripts not showing circadian behavior in our study, only 11 showed significant periodic behavior in the Wolff et al data. A Chi-square comparison (20 of 25 versus 11 of 25) gave a highly significant difference (P=0.0087). **Supplemental Figure 1 (**https://doi.org/10.25444/nhlbi.20533536) shows a graphical comparison of data from the two studies, showing highly similar time courses. Overall, the similarity of findings in the two studies provides important validation of the methods used in both.

### Limitations and future studies

This paper investigates circadian variation in mRNA levels in the proximal tubule. For protein-coding genes, mRNAs are translated to produce the proteins that fulfil their physiological functions. Circadian variation in mRNAs do not necessarily imply similar changes in the proteins they produce, however. For the abundance of a given protein to vary in response to circadian variations in their mRNAs, they must be degraded rapidly enough to fall when the mRNA abundance falls. Practically, half-lives must be less than a half cycle (<12 hours) for protein dynamics to match circadian mRNA dynamics. Prior studies have measured half-lives across the proteomes of living animals using metabolic labeling (65-68). We do not have comprehensive half-life data for native proximal tubule, but protein half-lives have been reported for native liver (67), which is metabolically similar to proximal tubule. In that study, of 1122 proteins quantified, the median half-life was 68.5 hours and only 56 proteins had half-lives that were less than 12 hours (5% of all proteins). Accordingly, we raise the question of what fraction of proteins vary in a circadian manner and whether more than about 5% of all proteins undergo circadian variation in native kidney proximal tubule. Direct metabolic labeling studies in mouse kidney seem warranted to address this question. Future studies also seem warranted to profile circadian changes in protein abundances.

## Supporting information

Supplemental Figure 1 (Comparison with Wolff et al.)

## DATA SHARING

All RNA-seq data can be downloaded from https://doi.org/10.25444/nhlbi.20533536.

## ACKNOWLEDGMENTS

The authors acknowledge expert help from the Bioinformatics Core Facility (NHBLI, Medhi Pirooznia, Director). M.A.B. was a member of the Biomedical Engineering Student Internship Program supported by the National Institute for Biomedical Imaging and Bioengineering (June-August, 2021, Robert Lutz, Director).

## AUTHOR CONTRIBUTIONS

M.D., K.N., and J.L. designed the experiments and conducted the experiments; M.A.B. and M.A.K. analyzed the data; M.A.B., M.A.K. and M.D. wrote the original version of the manuscript; H.K, H.J.J., B.G.P, V.R., E.P., C.R.Y., and C.L.C. reviewed the literature and provided advice and discussion of the data; all authors edited the manuscript and approved the final version.

